# Altered expression of MMP9 and TIMP1 leads to impaired placentation in early onset preeclamptic patients: a case control study

**DOI:** 10.1101/2023.03.08.531753

**Authors:** Pallavi Arora, Sankat Mochan, Sunil Kumar Gupta, Neerja Rani, Pallavi Kshetrapal, Sadanand Dwivedi, Neerja Bhatla, Renu Dhingra

## Abstract

**Introduction:** Matrix metalloproteinases (MMPs) specifically MMP-9 is a key regulator of vascular and uterine spiral artery remodelling and its activity is controlled at multiple levels, including gene transcription, activation of its latent forms and endogenous inhibition by tissue inhibitors of metalloproteinases [TIMPs (specifically TIMP-1)]. Alteration in MMP-9 and TIMP-1 expression may contribute to uterine and vascular dysfunction leading to adverse pregnancy outcomes such as Preeclampsia (PE). Depending on time, PE is classified as early-onset preeclampsia, which is diagnosed before 34 weeks of gestation, or late-onset preeclampsia, which is diagnosed after 34 weeks. Early-onset preeclampsia (EOPE) is a severe obstetrics disease which threatens mother and foetus. Therefore, in the present study, we aimed to investigate the levels of MMP-9 and its inhibitor TIMP-1 in the placentae of EOPE patients and their maternal age matched normotensive, non-proteinuric controls at both mRNA and protein levels.

**Methods:** A total of 30 caesarean delivered placentae (15 EOPE patients and 15 controls) were collected from Department of Obstetrics and Gynaecology, AIIMS, New Delhi after taking permission from Institute Ethics Committee. MMP-9 and TIMP-1 protein expression was observed by immunohistochemistry and immunofluorescence stainings. Gelatin Gel Zymography was done to evaluate MMP-9 activity and Western Blot was done for the quantification of MMP-9 and TIMP-1 proteins. mRNA levels of MMP-9 and TIMP-1 were determined by qRT-PCR

**Results:** Immunohistochemistry and Immunofluorescence staining discerned stronger expression of MMP-9 in normotensive placentae as compared to EOPE placentae whereas stronger expression of TIMP-1 was seen in EOPE placentae in comparison to those of normotensive placentae. Gelatin Gel Zymography reflected that MMP-9 activity was found elevated in maternal placental side in normotensive placentae as compared to EOPE placentae. Western Blot analysis revealed that protein expression of MMP-9 was elevated whereas for that of TIMP-1 was reduced in normotensive placentae in comparison to EOPE placentae. mRNA expression of MMP-9 was found up-regulated whereas for that of TIMP-1 was down-regulated in normotensive placentae as compared to EOPE placentae.

**Conclusion:** The present study implies that aberrant functioning of MMP-9 and TIMP-1 in EOPE patients contribute to impaired placentation which might be relevant for possible future screening programs in order to predict and to design therapies for early onset preeclamptic patients.

## Introduction

Normal pregnancy is associated with hemodynamic and uterine changes that allow adequate uteroplacental blood flow and uterine expansion for the growing fetus. These pregnancy associated changes require significant uteroplacental and vascular remodeling^1^. Matrix metalloproteinases (MMPs) are regulators of vascular and uterine remodeling^1^. Matrix metalloproteinase-9 (MMP-9) is the only effective hydrolase secreted by trophoblast cells, involved in extracellular matrix (ECM) remodeling and placental angiogenesis with structural spiral arteries transformation, which precedes proper trophoblastic invasion^2^. Tissue inhibitor of metalloproteinase-1 (TIMP-1), corresponding to MMP-9 inhibits latter’s activity to degrade collagen^3^. The balanced expression of MMP-9 and TIMP-1 has profound implications for regulating trophoblast invasion^3,4^. Increase in MMP-9 has been implicated in vasodilation, placentation, and uterine expansion during normal pregnancy^1^ however, reduced MMP-9 could impede uterine growth and expansion and lead to decreased vasodilation, hypertensive pregnancy, premature labour and preeclampsia (PE)^1^. Previous studies reported that for the specific inhibitor TIMP-1, no corresponding change was observed but the state of equilibrium between MMP-9 and TIMP-1 was altered in PE^3^. PE is a multisystem disorder of pregnancy defined by high blood pressure and proteinuria^5^ and has been classified into 2 different disease entities: early-onset preeclampsia (EOPE) and late-onset preeclampsia (LOPE)^6,7^. EOPE develops before 34 weeks of gestation, whereas late-onset PE develops at or after 34 weeks of gestation^6,7^. EOPE is associated with placental dysfunction, reduction in placental volume, intrauterine growth restriction, abnormal uterine and umbilical artery Doppler evaluation, low birth weight, multiorgan dysfunction, perinatal death, and adverse maternal and neonatal outcomes^6^. The results of previous research into the role and expression pattern of MMP9 and TIMP1 in pregnancies complicated by early onset preeclampsia are ambiguous. Therefore, the aim of this study was to explore whether placentae from early onset preeclamptic pregnancies were associated with altered expression of MMP-9 and TIMP-1 as compared to placentae from normotensive pregnancies.

## Methodology

### Sample size

15 each of early-onset preeclamptic (EOPE) patients and maternal age matched normotensive, non-proteinuric controls placentae (caesarean delivered) were collected from Department of Obstetrics and Gynaecology, AIIMS, New Delhi after taking ethical clearance from Institute Ethics Committee (IECPG-101/21.03.2018). EOPE patients were recruited as per ACOG (American College of Obstetrics and Gynaecology) guidelines. Patients with chronic hypertension, chorioamnionitis, diabetes, renal disease and cardiac disease were excluded from the study.

Protein expression of MMP-9 and TIMP-1 was observed by Immunohistochemistry (IHC) and Immunofluorescence (IF). Gelatin gel zymography was done to determine the proteolytic activity of MMP-9. Western Blot was carried out to determine MMP-9 and TIMP-1 protein levels. MMP-9 and TIMP-1 mRNA expressions were analysed using qRT-PCR. Relative normalized expression (Fold change) was calculated by 2^−ΔΔCt^ method.

### Immunohistochemistry

Paraffin tissue blocks were sectioned on microtome (Thermo Scientific™ HM 325) (5μm sections) and were taken on Poly-L-lysine (Sigma) coated slides. UltraVision™ Quanto Detection System HRP DAB (Thermo, TL-125-QHD) was used to determine MMP-9 and TIMP-1 protein expressions. Primary antibodies to MMP-9 (Abcam) at a dilution of 1:500 and TIMP 1 (Thermo) at a dilution of 1:100 were used. Slides were counterstained with hematoxylin, dehydrated in graded ethanol, cleared, and cover slips applied. Stained slides were observed under Nikon Eclipse Ti-S elements microscope using NiS-AR software (version 5.1). Other chemicals were procured from Fischer Scientific.

### Immunofluorescence

Paraffin tissue blocks were sectioned on microtome (Thermo Scientific™ HM 325) and were taken on Poly-L-lysine (Sigma) coated slides. Two changes of xylene (each for 5 minutes) followed by two changes of absolute alcohol (each for 3 minutes) and subsequently one change of 90% alcohol were given for 1 minute. Slides were rinsed with distilled water. Antigen retrieval was done with sodium citrate buffer (15 minutes at 95-100°C) followed by treatment with TBSTx (2 minutes) (2 times). BSA (blocking agent) was applied on slides for 35 minutes followed by overnight incubation with primary antibodies [MMP-9 (Abcam) at a dilution of 1:50 and TIMP 1 (Thermo) at a dilution of 1:10] at 4°C. Slides were then rinsed with PBST20 followed by incubation with secondary antibodies [MMP-9 (FITC conjugated, Abcam) at a dilution of 1:500 and TIMP 1 (TRITC conjugated, Thermo) at a dilution of 1:500] for 2.5 hours and then washed with PBS. Mounting was done with fluoroshield mounting media with DAPI (Abcam). Stained slides were observed under Nikon Eclipse Ti-S elements microscope using NiS-AR software (version 5.1). Other chemicals were procured from Fischer Scientific.

### Gelatin gel zymography

Protein extraction (from placental tissues of both patients and controls) was done with RIPA buffer (Thermo) and protease inhibitor cocktail. 7.5% acrylamide gel was prepared containing gelatin. 5x non-reducing sample buffer was added to the isolated protein samples. 10 μl protein sample was loaded to each well. Protein molecular weight marker (Thermo) was also loaded followed by running the gel at 90 V (vertical electrophoresis apparatus, BioRad) in electrophoresis buffer until good band separation is achieved. Gel was washed (2 × 30 min) with washing buffer then rinsing in incubation buffer for 5–10 min in incubation buffer at 37°C with agitation. Fresh incubation buffer was added to the gel followed by incubation for 24 h at 37°C. Gel was stained with Coomassie blue for 30 min and rinsed with water. Incubation was done with destaining solution to visualise the bands showing pro and active forms of MMP-9 in the study groups.

### Western Blot

Protein extraction (from placental tissues of both patients and controls) was done with RIPA buffer (Thermo) and protease inhibitor cocktail. Separating and stacking gels were prepared. 3x non-reducing sample buffer was added to the isolated protein samples followed by denaturation at 95^°^C for 5 minutes and the samples were loaded along with protein molecular marker to the wells. Subsequently, running the gel at 50 V (vertical electrophoresis apparatus, BioRad) in electrophoresis buffer for 3-3.5 hours until good band separation is achieved. Nitrocellulose membrane was used for the transfer of gel products on to the membrane in transfer buffer for 90 minutes. Membrane was washed with TBST20 followed by blocking in 5% BSA TBST20 for 90 minutes. Overnight incubation was done with primary antibodies [MMP-9 (Abcam) at a dilution of 1:1000 and TIMP 1 (Thermo) at a dilution of 1:500] at 4°C. Washing was done with TBST20 followed by incubation with secondary antibodies [MMP-9 (Abcam) at a dilution of 1:1000 and TIMP 1 (Thermo) at a dilution of 1:1000] for 3 hours and then washed with TBST20. ECL kit (Thermo) was used for visualization of bands in Densitometer (Protein Simple).

### qRT-PCR

Placental tissue (kept in RNA later) was blotted on absorbent paper and 100 mg was taken, washed in 1x 0.1 M PBS. RNA isolation was done using Ambion, Invitrogen. The quality of RNA was examined by denaturing gel and quantity was measured on Micro-Volume UV/Visible Spectrophotometer (Thermo Fisher Scientific-NanoDrop TM 2000). c-DNA synthesis was done using Thermo revert aid H-minus reverse transcriptase kit (Thermo). Quality of cDNA was checked on 0.8% agarose gel visualized by ethidium bromide (EtBr) stain under UV and quantity was measured on Micro-Volume UV/Visible Spectrophotometer. cDNA was subsequently used for qRT-PCR. qRT-PCR reactions were carried out in 20 μl volume, including SYBR Green (Thermo), forward and reverse primers [MMP-9, TIMP-1 (Sigma)], cDNA (template) and nuclease free water (CFX96 Touch™ Real-Time PCR Detection System, BioRad). Initial Denaturation was performed at 95°C for 3 minutes, final at 95°C for 15 seconds, annealing at 60°C and extension at 70-72° C. Primers were designed by NCBI and confirmed by *In silico* PCR (Table 1). GAPDH was used as reference gene.

**Table 1:**
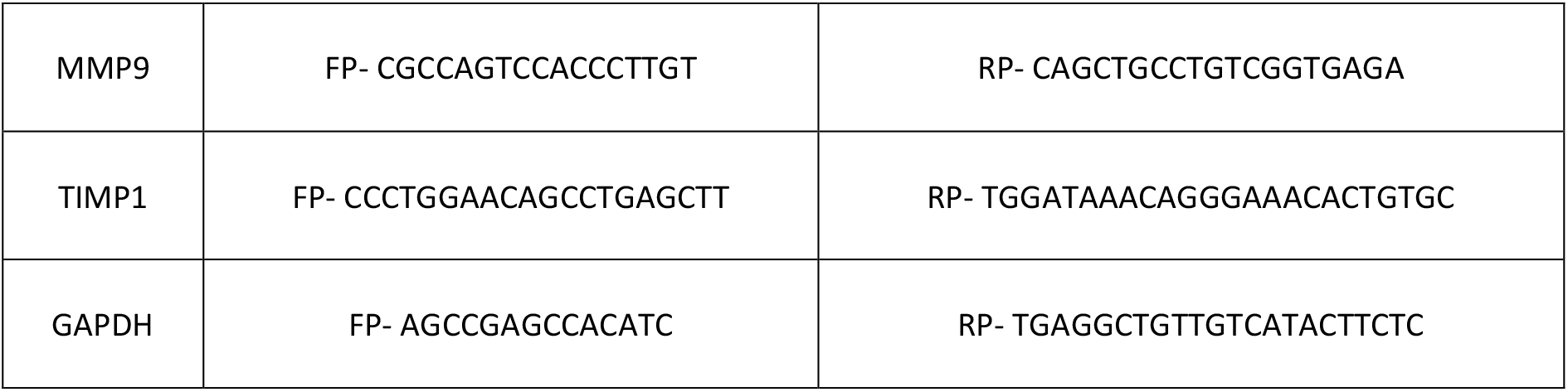
Primers are designed by NCBI (National Centre for Biotechnology Information) and confirmed by *In silico* PCR. GAPDH: Glyceraldehyde 3-phosphate dehydrogenase

### Statistical Analysis

Data was analyzed by STATA 14.1 and GraphPad Prism 9.4.1. For qRT-PCR data analysis, relative quantification cycles of gene of interest (ΔCq) was calculated by ΔCq = Cq (target) - Cq (reference). Relative normalized expression (Fold change) was calculated by 2^-ΔΔCq^ method. *p* value<0.05 was considered statistically significant.

## Results

Clinical characteristics of early onset preeclamptic patients and maternal age matched normotensive, non-proteinuric controls are mentioned in Table 2.

**Table 2:** Maternal study population - clinical characteristics. Data represented as mean± SD. *p* values between groups are evaluated by Paired t-test. *p*<0.05 considered statistically significant.

| Clinical features | EOPE patients (n=15) | Controls (n=15) | *p* value (Paired-t test) |
| --- | --- | --- | --- |
| Maternal Age (years) | 29.9 ± 4.09 | 30.1 ± 3.69 | 0.76 |
| Systolic BP (mm Hg) | 162.1 ± 11.58 | 117.5 ± 6.32 | <0.0001 |
| Diastolic BP (mm Hg) | 101.4 ± 9.25 | 76.1 ± 6.27 | <0.0001 |
| Proteinuria (Dipstick) | 3+ = 5 (33.33%), 2+ = 7 (46.66%), 1+ = 3 (20%) | nil or in traces | Not applicable |
| Placental weight | 440 ± 47.71 | 507.8 ± 9.60 | 0.04 |
| Intrauterine Growth Restriction (IUGR) | 3 (20%) | 0 (0%) | Not applicable |

### Weak expression of MMP9 and enhanced signalling of TIMP 1 in EOPE placentae as compared to normotensive, non-proteinuric (controls) placentae was observed (Figure 1-4)

The expression of MMP-9 was reduced in early onset preeclamptic(EOPE) placentae as compared to placentae from normotensive pregnancies as seen in both immunohistochemistry and immunofluorescence staining (Figures 1a-b, 3e, 3h 3b). However, intense immunostaining for TIMP1 was observed in both the syncytium and stromal compartment in EOPE placentae (Figures 2a, 2c, 4b, 4e) as compared to control placentae (Figures 2b, 2d, 4h).

**Figure 1:**
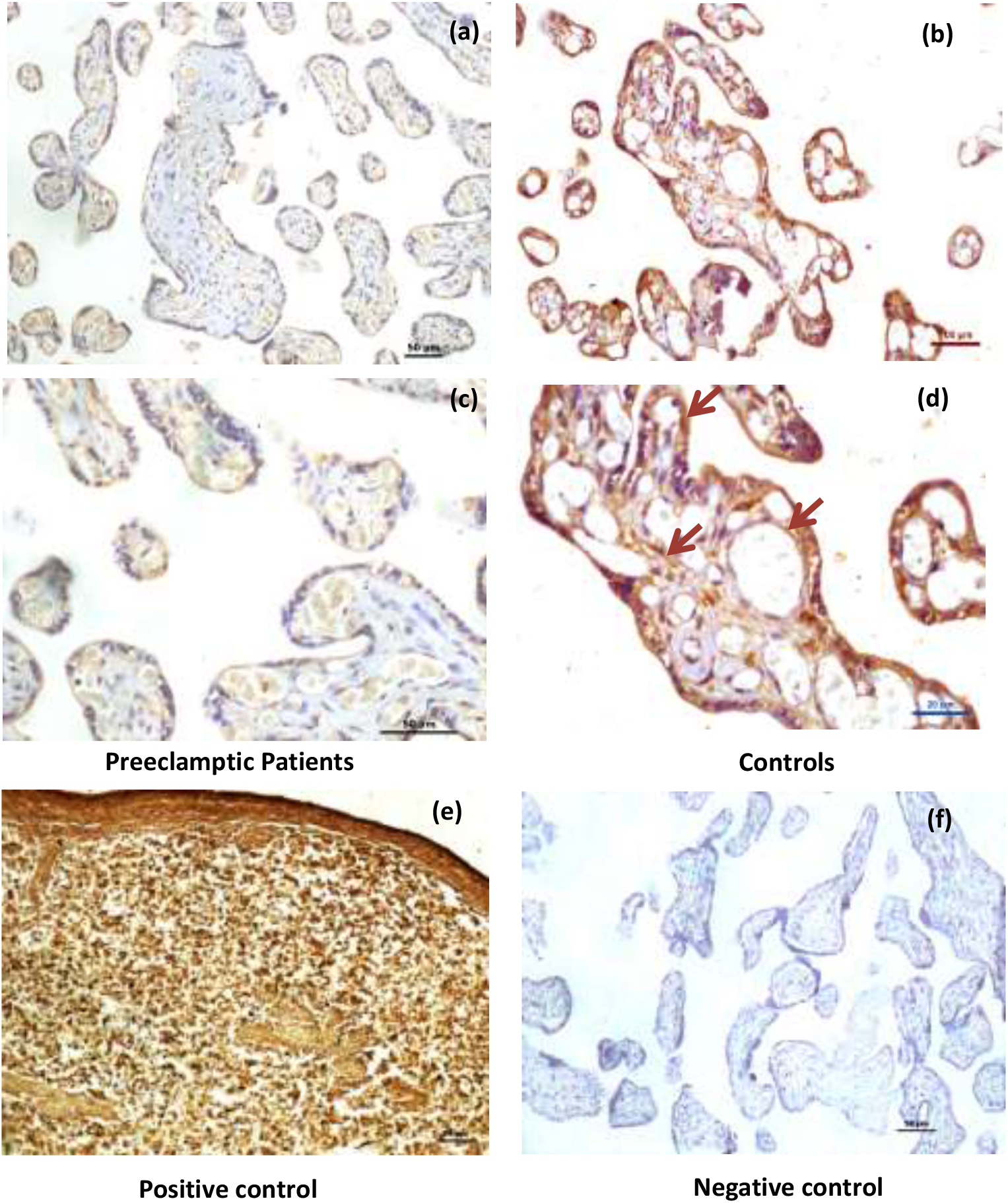
Representative Immunohistochemistry images of placentae from EOPE patients (a: 20x, c: 40x) and normotensive, non-proteinuric controls (b: 20x, d: 40x) showing MMP-9 localization. Arrows refer to MMP-9 positive cells. Human spleen serves as positive control for MMP-9 expression (e). (f): negative control.

**Figure 2:**
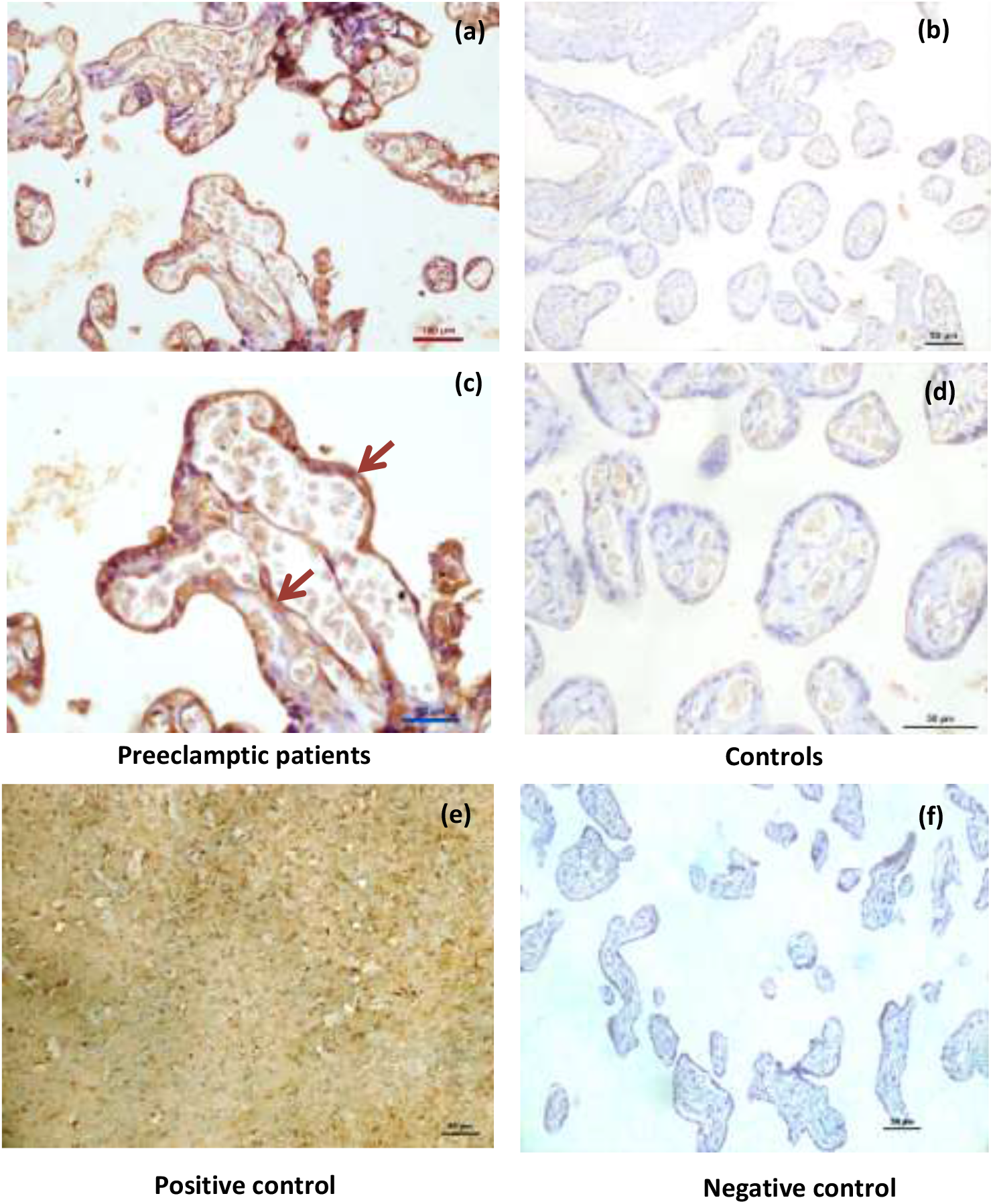
Representative Immunohistochemistry images of placentae from EOPE patients (a: 20x, c: 40x) and normotensive, non-proteinuric controls (b: 20x, d: 40x) showing TIMP-1 localization. Arrows refer to TIMP-1 positive cells. Human brain tissue serves as positive control for TIMP-1 expression (e). (f): negative control.

**Figure 3:**
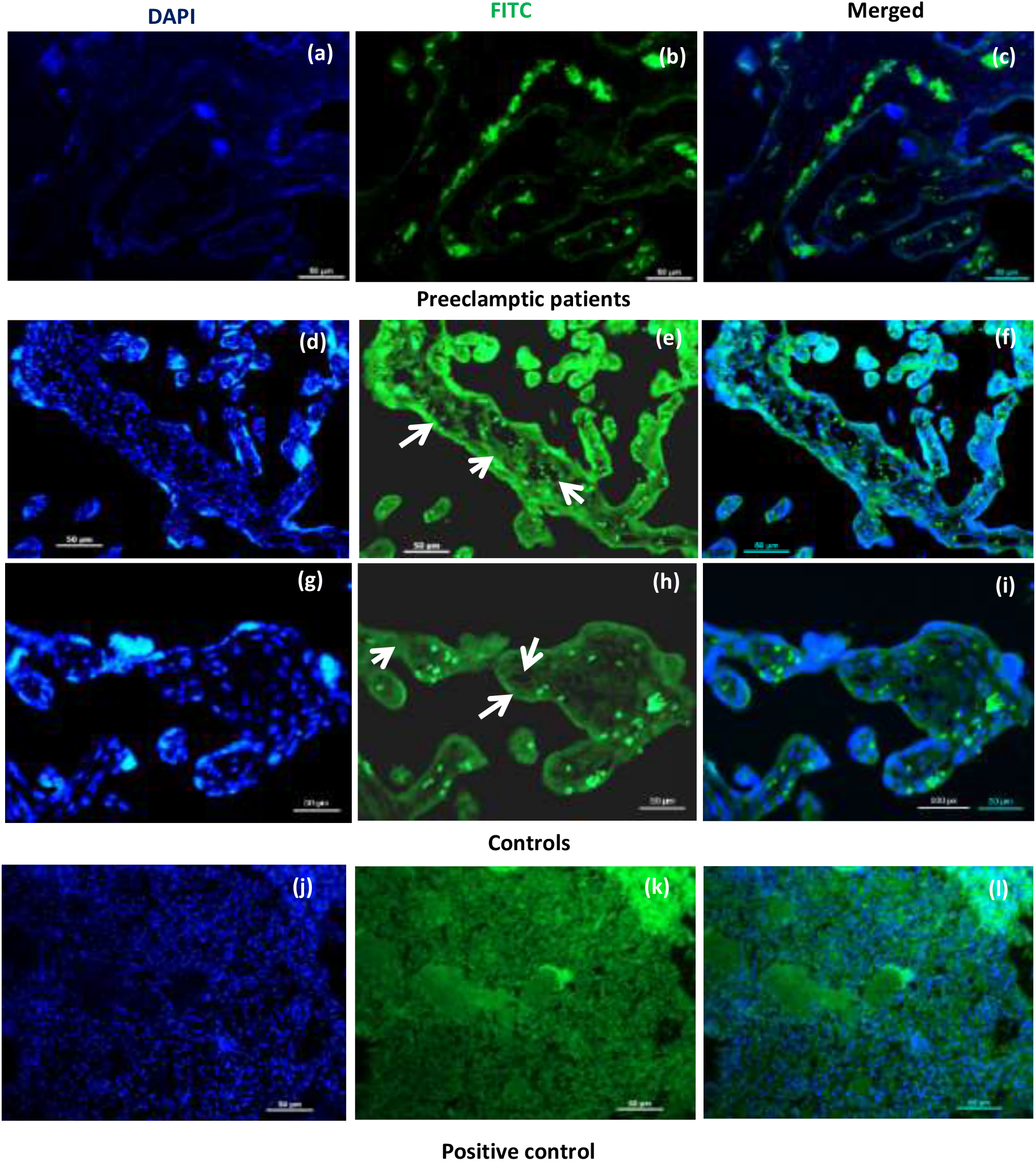
Representative immunofluorescence images of placentae from EOPE patients (a-c) and normotensive, non-proteinuric controls (d-i) showing MMP-9 localization [FITC]. Nuclei are stained by DAPI. Arrows refer to MMP-9 positive cells. Human spleen serves as positive control for MMP-9 expression (j-l).

**Figure 4:**
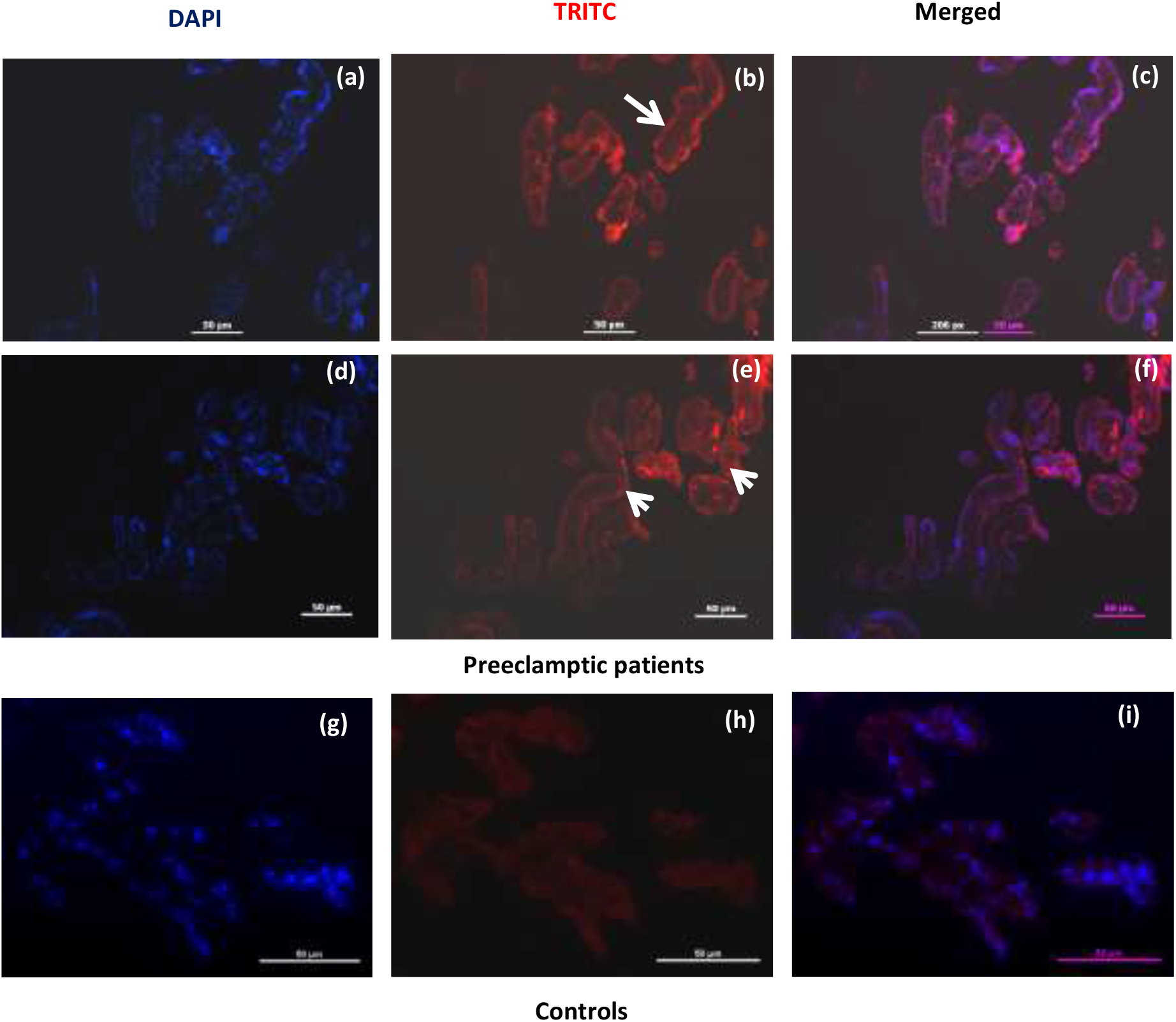
Representative immunofluorescence images of placentae from EOPE patients (a-f) and normotensive, non-proteinuric controls (g-i) showing TIMP-1 localization [TRITC]. Nuclei are stained by DAPI. Arrows refer to TIMP-1 positive cells.

### Gelatinolytic activity of MMP-9 was reduced in early onset preeclamptic patients

MMP-9 activity [both pro (*p*=0.02) and active forms (*p*=0.04)] was significantly lower in placentae from early onset preeclamptic patients (Figures 5a, 5c, 5d) as compared to normotensive, non-proteinuric controls (Figures 5b, 5c, 5d).

**Figure 5:**
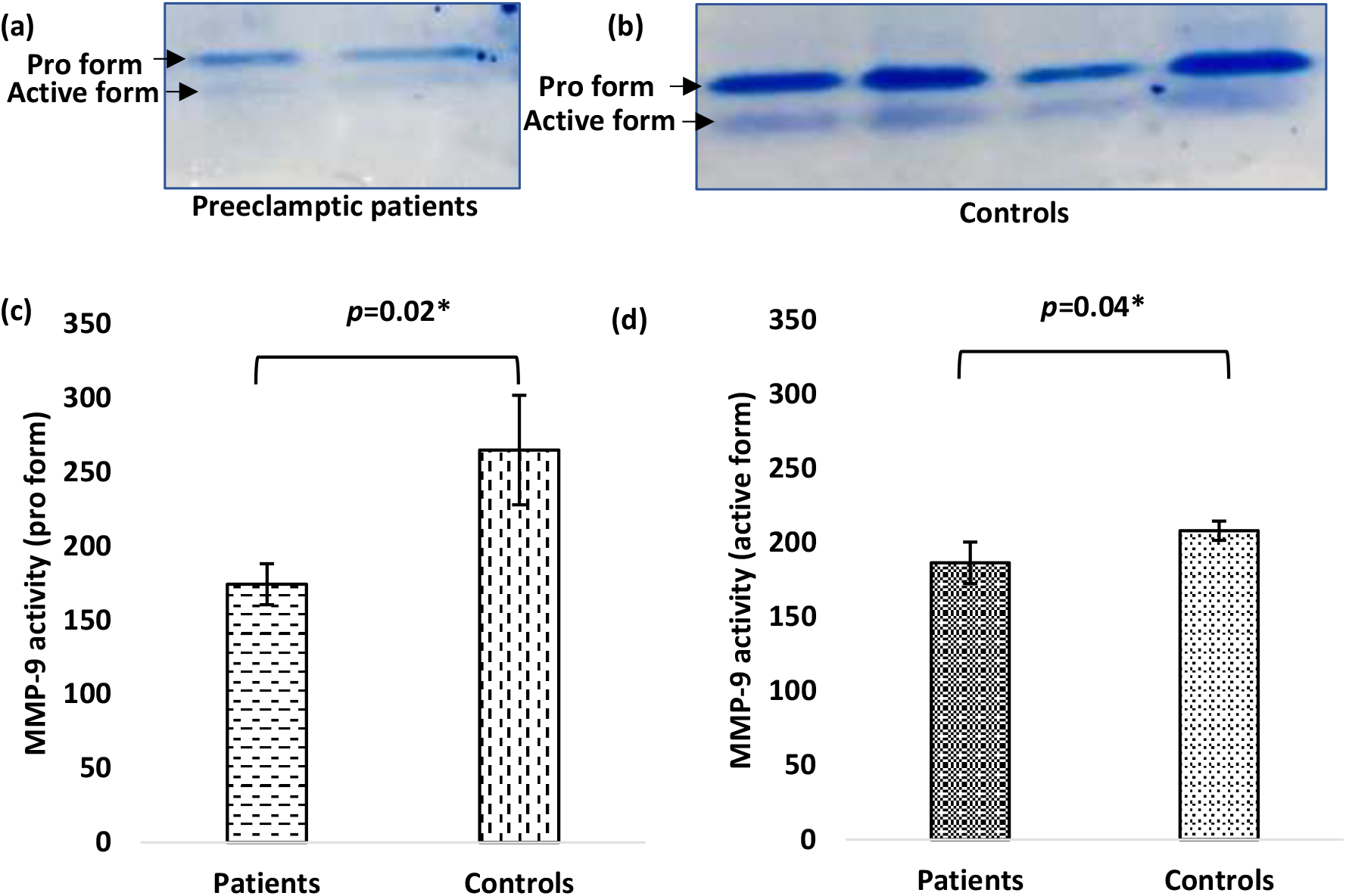
Representative zymograms depicting gelatinase activity on maternal side of placentae from EOPE patients (a) and normotensive, non-proteinuric controls (b). Comparison of Pro (92 KDa) (c) and active (86 KDa) (d) forms of MMP-9 between patients and controls by Paired t-test. Data presented as mean ± SD. *p*<0.05 considered statistically significant.

### Reduced MMP-9 and elevated TIMP-1 protein expressions in early onset preeclamptic patients

Immunoblot data revealed significant decrease (*p*=0.04) in MMP-9 (Figures 6a, 6c) and increase (*p*=0.01) for that of TIMP-1 (Figure 6b, 6d) protein expression in placentae from early onset preeclamptic patients as compared to normotensive, non-proteinuric controls.

**Figure 6:**
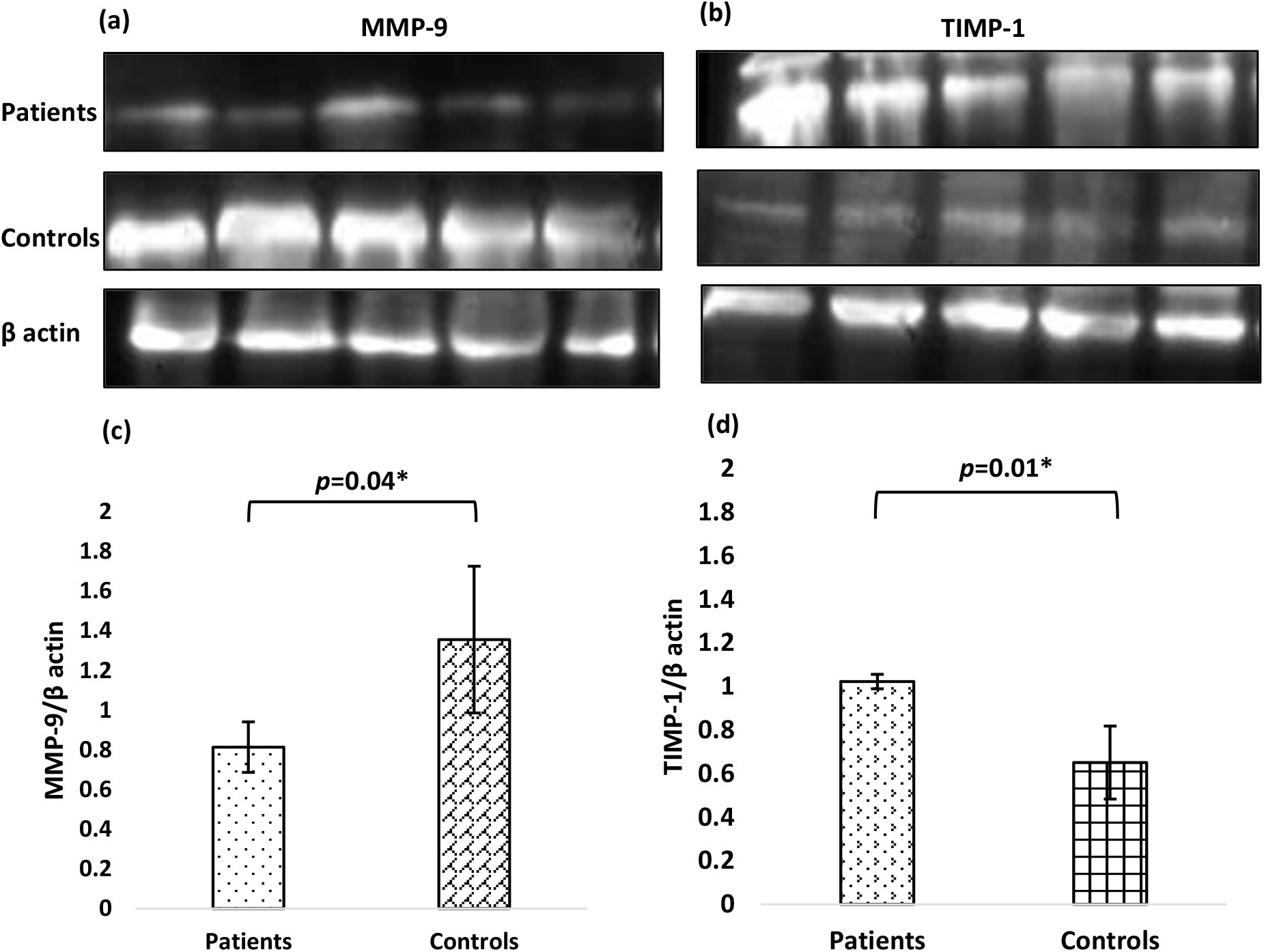
Immunoblots showing MMP-9 (a) and TIMP-1 (b) protein expressions in placental tissues of EOPE patients and normotensive, non-proteinuric controls. β actin: protein loading control. Comparison of normalized protein expression of MMP-9 (c) and TIMP-1 (d) between the two groups (Patients and Controls) by Paired t-test. Data presented as mean ± SD. *p*<0.05 considered statistically significant.

### Downregulated gene expression of MMP-9 whereas upregulated for that of TIMP-1 in early onset preeclamptic patients was seen

qRT-PCR data showed 6.28 folds reduction of the mRNA expression of MMP-9 (Figure 7a) in placentae from early onset preeclamptic patients as compared to that of normotensive non-proteinuric controls whereas 8.32 folds increase in the mRNA expression of TIMP 1 (Figure 7b) was observed in placentae from early onset preeclamptic patients.

**Figure 7:**
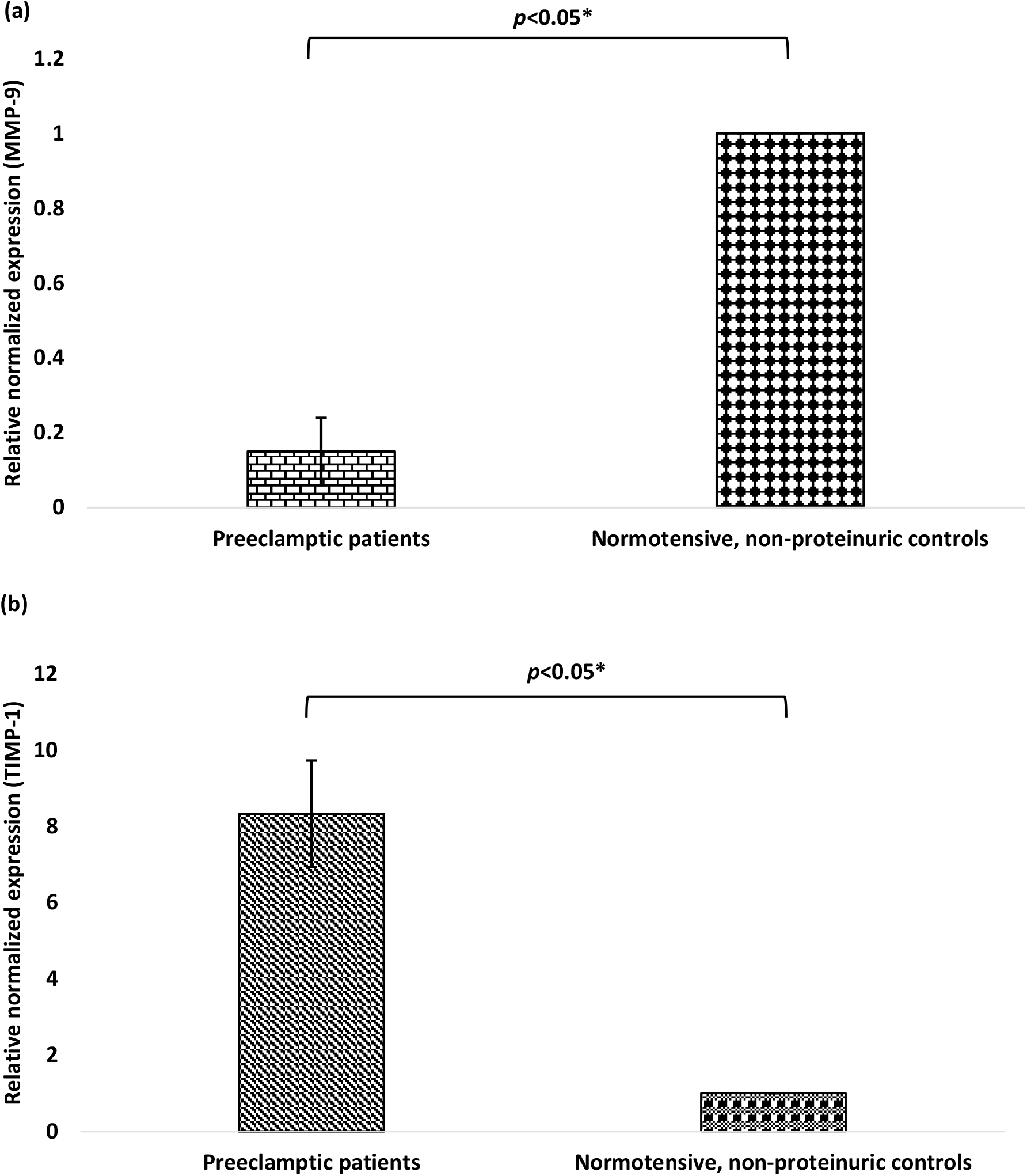
Relative normalized expression (Fold change) was calculated and compared for MMP-9 (a) and TIMP-1 (b) genes between EOPE patients and normotensive, non-proteinuric controls by 2-^ΔΔCt^ method.

## Discussion

In the present study, we have analysed the both qualitative and quantitative protein and mRNA expression of MMP-9 and TIMP-1 in 15 pairs of placentae from EOPE patients and their maternal age matched normotensive, non-proteinuric controls.

Matrix metalloproteinases (MMPs) are essential biochemical mediators to facilitate trophoblast cell’s invasion and adequate spiral artery remodelling^8,9^. MMPs, also called matrixins, are a family of 17 zinc-dependent endopeptidases, which participate in many biological processes^8,9^. The regulation of MMPs activity at the maternal-fetal interface appears to be critical for successful placentation^8,9^. Matrix metalloproteinase-9 (MMP-9) is the only effective hydrolase secreted by the trophoblast cells which can digest ECM proteins^3^. MMP-9 is strongly localized in the placental bed (primarily to EVCTs) and appears to regulate trophoblast invasion^10^. Appropriate trophoblast invasion and vascularization requires a functional synergism between MMP-9 and its regulating factor TIMP-1^3^.

We observed reduced expression of MMP-9 in syncytiotrophoblasts, stroma and around the blood vessels in placentae from early onset preeclamptic women as compared to placentae from normal pregnant women. The reduced expression of MMP-9 in trophoblast cells could cause shallow invasion, disturbed and inadequate remodelling of the maternal spiral arteries by invading trophoblast cells, thus reducing blood flow to the intervillous space^3,11^. Insufficient conversion of the spiral arteries into low-resistance, high capacity vessels in early pregnancy can lead to systemic hypertension and fetal hypoxia in later pregnancy as the fetus and placenta outgrow their blood supply, often observed in PE especially the early onset type (EOPE)^11^.

EOPE is the most severe clinical variant of disease occurring in 5-20% of all cases of PE and is associated with impaired fetal growth, fetal pathology and uterine blood circulation, small size of the placenta, preterm delivery, neonatal morbidity and mortality^12^. EOPE developments are associated with impaired trophoblast invasion, immune maladaptation and increased markers of endothelial dysfunction^13^. However Late Onset type (LOPE) is occurring in about 75-80% of all cases of preeclampsia, associated with maternal morbidity (metabolic syndrome, impaired glucose tolerance, obesity, dyslipidemia, chronic hypertension) but normal birth weight and normal placental volume^14^.

We have observed strong signalling of TIMP-1 in the syncytium and stroma in EOPE placentae. Our results corroborate with the previous study by Zhang *et al*., 2019 where they observed enhanced intensity of MMP-9 in cytoplasm of trophoblast cells, vascular endothelial cells and villous mesenchymal cells in healthy placentae^3^. However, they also observed that the specific inhibitor of MMP-9 i.e. TIMP-1 had no corresponding change and the equilibrium state between MMP-9 and TIMP-1 changed^3^.

Another study proposed that MMP-9 ablation in MMP-9 knockout mice shows a phenotype that mimics PE due to impaired trophoblast differentiation and invasion^15^. Shokry *et al*. in 2009 and Omran *et al*. in 2011 both reported decreased MMP-9 levels in preeclamptic patients as compared to controls^16,17^. Dang *et al*. in 2013 confirmed that MMP-9 was abundantly expressed in tissues of normal pregnant rats by immunohistochemistry, gelatin gel zymography and Western blot analysis supporting a role of MMP-9 in the uteroplacental and vascular remodelling during normal pregnancy^18^.

In the present study gelatin gel zymography, revealed that both the pro and active forms of MMP-9 were significantly reduced in placentae from EOPE patients as compared to their maternal age matched normotensive, non-proteinuric controls The quantitative estimation of MMP-9 and TIMP-1 proteins by western blot found significant decrease in the levels of MMP-9 and increase for that of TIMP-1 in EOPE placentae thereby validating the immunostaining (immunohistochemistry and immunofluorescence) results. Gene expression pattern of MMP-9 and TIMP-1 was also altered in EOPE patients.

Aberrant expression of MMP-9 and TIMP-1 has been implicated in pregnancy abnormalities, including IUGR and preeclampsia (PE)^3^. Moreover, excessively shallow invasion has been implicated more in EOPE and not as much in LOPE^8,9^.

The molecular mechanism behind the development of EOPE is unexplored. Therefore, in the present study, we hypothesized that the altered levels of matrix metalloproteinase-9 and its regulator, tissue inhibitor of metalloproteinase-1 may contribute to the pathogenesis of EOPE.

## Conclusion

Reduced expressions of MMP-9 and increased expression of TIMP-1 were observed in placentae from early onset preeclamptic patients as compared to normotensive, non-proteinuric controls at both protein and mRNA levels. We have shown altered protein expression of these two markers not only at cellular level but also the altered enzyme activity in EOPE placentae. The results were also validated by immunoblot. The significant changes in the protein expression were also correlated with mRNA expression in placentae of these patients. The role of MMP-9 and TIMP-1 as key mediators of pathogenesis of early onset preeclampsia needs to be further explored. Longitudinal prospective studies can be designed to analyze their use as potential biomarkers and biological therapeutic targets.

## Notes

### Competing Interest Statement

The authors have declared no competing interest.

